# zDB: bacterial comparative genomics made easy

**DOI:** 10.1101/2023.05.31.543076

**Authors:** Bastian Marquis, Trestan Pillonel, Alessia Carrara, Claire Bertelli

## Abstract

The analysis and comparison of genomes relies on different tools for tasks such as annotation, orthology prediction and phylogenetic inference. Most tools are specialized for a single task and additional efforts are necessary to integrate and visualize the results. To fill this gap, we developed zDB, an application that integrates an analysis pipeline and a visualization platform. Starting from annotated Genbank files, zDB identifies orthologs and infers a phylogeny for each orthogroup. A species phylogeny is also constructed from shared single-copy orthologs. The results can be enriched with Pfam protein domain prediction, COG and KEGG annotations and Swissprot homologs. The web application allows searching for specific genes or annotations, running Blast queries and comparing genomic regions and whole genomes. The metabolic capacities of organisms can be compared at either the module or pathway levels. Finally, users can run queries to examine the conservation of specific genes or annotations across a chosen subset of genomes and display the results as a list of genes, Venn diagram or heatmaps. Those features will make zDB useful for both bioinformaticians and researchers more accustomed to laboratory research. zDB is perfectly suited to process datasets with tens to hundred of genomes on a desktop machine.

**IMPORTANCE:** Genome comparison and analysis rely on many independent tools, leaving to scientists the burden to integrate and visualize their results for interpretation. To alleviate this burden, we have built zDB, a comparative genomics tool that includes both an analysis pipeline and a visualization platform. The analysis pipeline automates gene annotation, orthology prediction and phylogenetic inference, while the visualization platform allows scientists to easily explore the results in a web browser. Among other features, the interface allows users to visually compare whole genomes and targeted regions, assess the conservation of genes or metabolic pathways, perform Blast searches or look for specific annotations. Altogether, this tool will be useful for a broad range of applications in comparative studies between two to hundred genomes. Furthermore, it is designed to allow sharing datasets easily at local or international scale, thereby supporting exploratory analyses for non-bioinformaticians on the genome of their favorite organisms.

## INTRODUCTION

Since the publication of the first complete genome in 1995 (1), the number of available sequences has kept on growing, with now 450’000 different species available in the Genbank database (2). As recent sequencing technologies make it possible to sequence an organism in a matter of hours at a cost affordable even for small research laboratories, this trend is unlikely to abate in the future. These technological improvements transferred the burden from sequencing to the actual analysis of the sequences. While a plethora of different tools already exist for this purpose, they are often specialized for specific tasks such as gene calling, orthology prediction or phylogenetic inference. Moreover, these tools are often standalone programs that do not readily integrate each other’s results. As the results are often not produced in a format that easily allows their exploration, additional visualization efforts are also necessary.

The need for tools designed to aggregate results from different sources has been illustrated by the success of programs like Prokka (3), which merges the results of different annotation tools in files ready for submission and visualization in genome browsers. The idea is further extended by pipelines like Bactopia (4), TORMES (5) and ASA3P (6) that automate all steps from reads quality control to antibiotic resistance gene prediction and generate simple HTML reports allowing the visualization of the main results. As these pipelines were developed with a focus on clinical microbiology, they are limited in terms of comparative genomics analysis. In contrast, websites dedicated to the comparative genomics of specific group of organisms (7, 8) have been developed and implement powerful interfaces allowing users to make custom queries and to generate complex plots. However, these websites do not allow users to analyze their own sets of genomes. Some web-based comparative genomics platforms, like EDGAR (9), PhyloCloud (10), KBase (11), CoGe (12) or MicroScope (13) implement similar interfaces while allowing users to upload their own dataset. Some of those platforms are however closed-source, and as the analysis are performed on the platform’s respective servers, users are required to register and upload their dataset. The ideal comparative genomics platform would therefore be open source, could run on any infrastructure, be as flexible and scalable as Bactopia, and similarly to MicroScope and EDGAR, offer an extensive interface to visualize the results. The developers of anvi’o (14) and bioBakery (15), both of which implement an analysis pipeline and a visualization platform demonstrate the feasibility of such an approach. bioBakery was however designed for microbial community profiling, and while anvi’o includes some analyses commonly performed in comparative genomics, such as pangenome analysis, it is not the main focus of this tool and some features available in EDGAR or MicroScope are lacking.

To fill this gap, we developed zDB, an open-source comparative genomics analysis pipeline and visualization platform. The analysis pipeline performs functional annotations, orthology and phylogenetic inference, while the visualization platform offers an interactive modern web-based interface to explore the results. Altogether, the ease of installing and executing the tool and the ability to easily visualize the results will benefit both bioinformaticians and researchers more accustomed to lab work.

## RESULTS

The visualization platform can be started by a single command as soon as the analyses are complete. The command starts a web server that will make the results available via a web browser, either only locally or also possibly extended to the whole Internet depending on the setup. The platform can also be used to visualize archived results imported from another computer.

### Visualization toolkit

The visualization platforms implements a set of plots and queries to explore the results of orthology prediction and phylogeny inference. In addition, zDB comes with several features of more general interest like the possibility to run Blast queries, to search for specific annotation or gene using a search bar and to draw interactive Circos plots or genomic regions.

A side bar is present on all pages of the web application (Figure 1A) and allow a quick access to all available analyses. The content of the “Annotations” tab varies in function of which optional analyses were performed. Similarly, the “Metabolism” tab will only be present when the genomes were annotated with KEGG orthologs. Summaries of the main characteristics of the genomes of interest, including the results from CheckM, can be visualized either as lists or directly annotated in the species phylogeny (Supplementary Fig. 1) via the “Genomes” and “Phylogeny” tabs, respectively. Finally, the web interface also includes a search bar (Figure 1A) that allows users to look for genes, gene products, bacteria or specific annotations based on their name. The search bar accepts wildcards and logical operators to combine different search terms.

**FIG 1.**
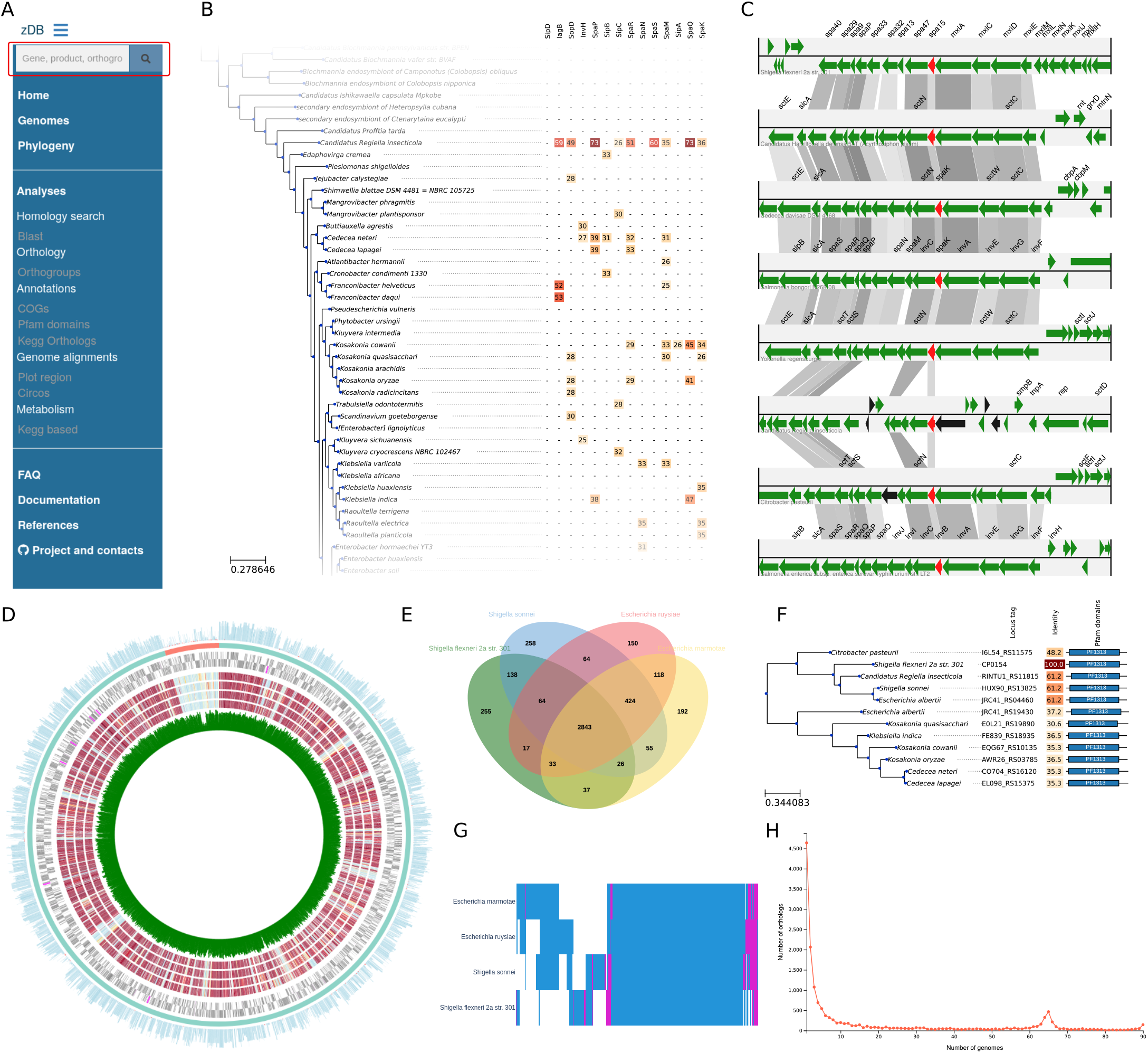
(A) zDB side panel listing all available analyses. The search bar (red box) allows users to look for a gene or annotation of interest. (B) Blast search result. The whole dataset was searched for several proteins of the type III secretion system of *Salmonella* Typhimurium. Blast hits are displayed as a heatmap of identities linked to the species phylogeny (cropped in this figure). (C) zDB can draw a comparison of the genomic region of a specified gene of interest and its orthologs (in red) in several genomes. Black arrows represent pseudogenes. Gray bars link orthologous genes, and their shade reflect amino-acid identity. (D) Visualization of protein conservation of a reference genome (here *Shigella flexneri*) in a set of 4 selected genomes (here from *Escherichia* genus). The inner circle represents the GC content of each open reading frame (ORF) in the reference genome. The next four circles represent the absence (in blue) or presence (in red) of homologs of proteins from the reference genome in the selected genomes, with a color scale representing protein identity. The next two circles represent the localisation of the ORFs on the forward and reverse strands of the reference genome. The two outer circles represent the contigs in the reference assembly (here two contigs: the chromosome and a plasmid) and a histogram of the number of homologs to each protein of the reference genome in the selected genomes. (E) Venn diagram illustrating the distribution of the orthologous groups in 4 genomes. (F) Gene phylogeny. The identity column shows the identity relative to the CP0154 locus, as it was accessed through the page dedicated to this locus. The rightmost columns displays the domain architecture of homologs (only present if the Pfam analysis is performed). (G) Heatmap of gene conservation in the 4 genomes as in (E); pink bars represent genes present in multiple copies in a genome, blue bars represent single copy genes. (H) Distribution of the orthologous groups in function of the number of genomes where they occur.

The “Orthology” tab links to pages allowing to explore gene conservation in the dataset. In particular, users can visualize gene conservation across a chosen set of genomes as either heatmaps (Figure 1G), Venn diagrams (Figure 1E) or lists. zDB can also draw the commonly used core and pan-genome plots as well as a plot of the number of orthogroups in function of the number of genomes where they occur (Figure 1H). The latter plot allows to quickly assess the number of singletons, the size of the core genome and to detect group of genes occurring in a subset of the genomes. Finally, zDB implements an interface that allows users to search for genes conserved in a chosen set of genomes but absent in another one.

As searching for specific sequences in organisms of interest is a frequent task, the visualization platform implements an interface that allows users to run their own Blast searches on either the whole dataset or on a specific genome. Several types of blast searches can be performed (blastp, blastn and tblastn), either with a single query or with multiple query in FASTA format. The results are displayed interactively using the BlasterJS library (16). Moreover, if the search was run on the whole dataset, zDB can display the results as a heatmap of the best blast hits identity linked to the species phylogenetic tree (Figure 1B). This allows to quickly detect patterns in the distribution of Blast hits in function of the phylogenetic distance.

Finally, zDB can draw plots to compare genomic regions sharing orthologous genes (Figure 1C) and Circos plots to compare a set of genomes to a specified reference (Figure 1D). The minimal setup also includes summary pages for every gene and orthogroup. The gene summary page allows to easily access the nucleotide and aminoacid sequences (for protein coding genes), displays the genomic region of the gene of interest, as well as the list of orthologous genes (Figure 2A). The orthogroup page allows users to examine the gene phylogeny and the distribution of the orthogroup in the genomes of the dataset (Figure 1F and Figure 2C). Both pages are enriched with the results of the optional analyses if they were performed (Figure 2A and 1F).

**FIG 2.**
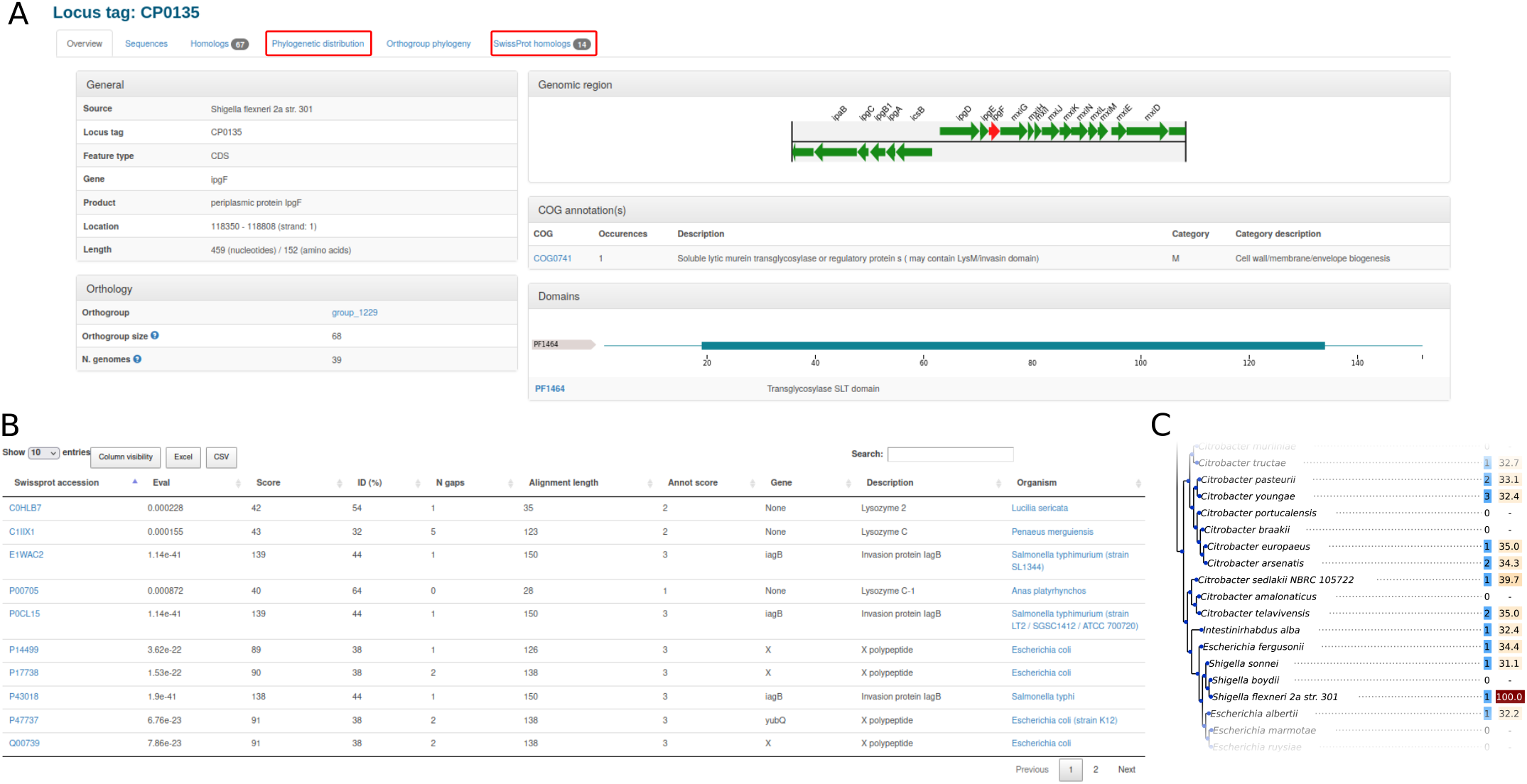
(A) Example of a gene summary page, with its genomic region, Pfam domains and COG annotation. The phylogenetic distribution and list of Swissprot homologs shown in (B) and (C) can be accessed from the highlighted tab. (B) The list of Swissprot homologs. (C) Part of the phylogenetic tree and protein conservation. The first column shows the number of homologs of the gene in a given genome. The second column shows the amino-acid identity between the gene of interest and its closest homolog in a given genome. 10.1128/aem.00309-23

### Pfam, COG and KEGG functional analyses

The conservation of Pfam, COG and KEGG annotations across genomes can be compared in a similar way to orthogroups. In particular, Venn diagrams, heatmaps, pan- and core-genome plots can be drawn for those annotations, while an interface to search for annotations present in a set of chosen genomes but absent in another is also available. Since COG and KEGG orthologs are assigned to high-level functional categories, users can visualize the distribution of annotated genes in those categories across one or several genomes, either as barcharts (Figure 3A and B) or as heatmaps (Figure 3C). This allows users to quickly visualize differences of functional capabilities between organisms.

**FIG 3.**
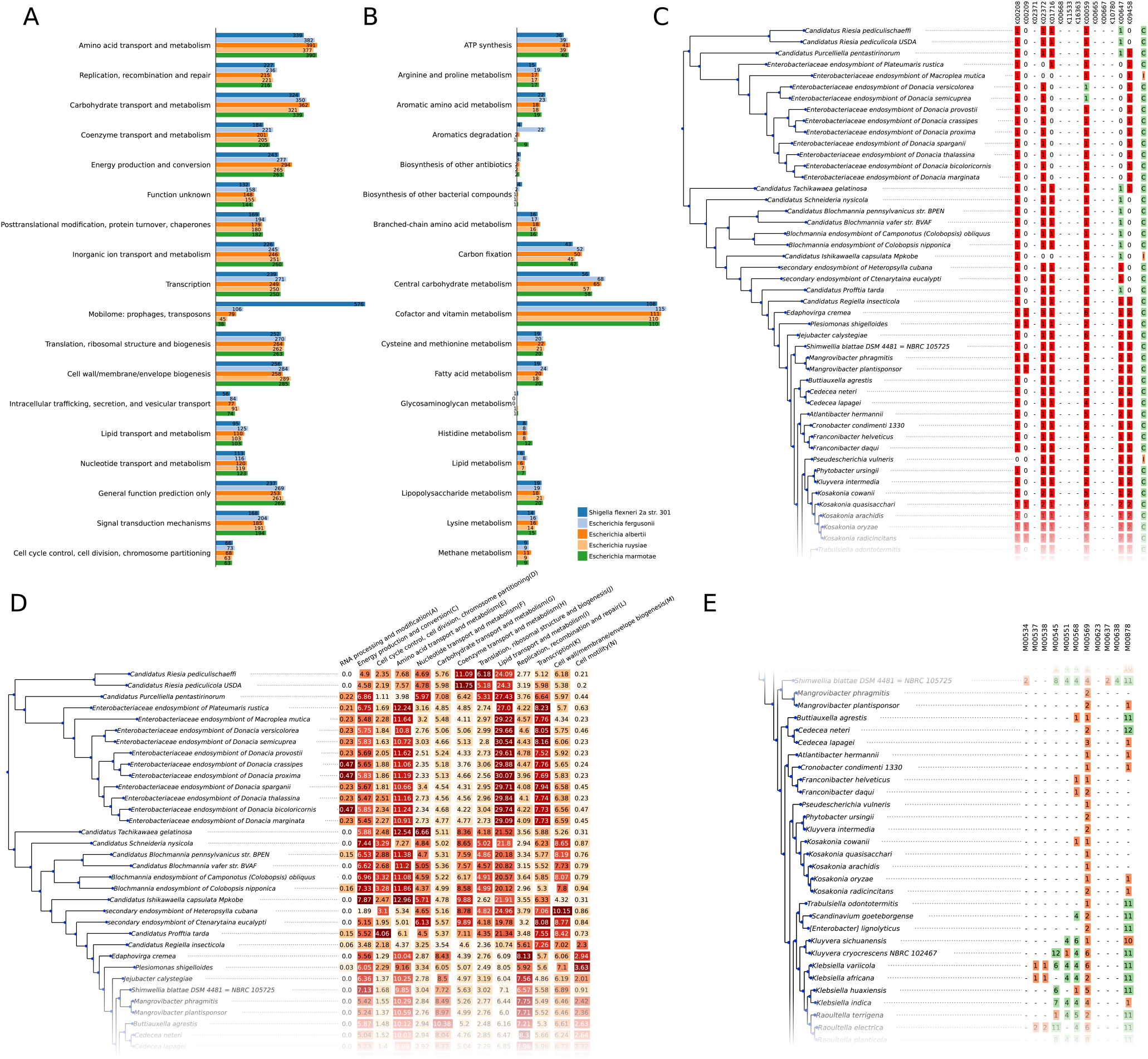
(A) and (B) Distribution of genes annotated with COG and KEGG orthologs in their functional categories for 4 chosen genomes. (C) Details of the completeness of KEGG module 83 (fatty acid biosynthesis, elongation). Red squares in the heatmap correspond to the number of genes annotated as a given KEGG ortholog. Green squares correspond to a gene without a specific KEGG annotation but in the same orthogroup as other genes having this annotation. This may indicate a shared function and in such cases, the corresponding KEGG ortholog is considered as present when estimating the module completeness. The last column indicates module completeness, as determined by the module definition language. (D) Proportion of the genes in a genomes assigned to the different COG categories (some categories were removed for the sake of simplicity). (E) Overview of modules completeness in any given KEGG category. Green squares indicate a complete KEGG module, orange squares indicate an incomplete module. The number of genes annotated as KEGG orthologs for a given module are indicated in each square.

To further characterize metabolic capacities, zDB implements a parser for the KEGG module definition language, which allows to assess the completeness of a metabolic module based on the KEGG orthologs present in a genome. Module complete-ness can be compared at the scope of a single KEGG module (Fig Figure 3C), or at the scope of categories or sub-categories (Fig Figure 3E). The results are directly linked to the species phylogeny, making it easy to notice patterns of metabolic capacities linked to specific clades.

The Swissprot homologs are listed both in the orthogroup home page and in the gene homepage (Fig Figure 2B).

### Benchmarking

An initial benchmark evaluated the duration of computations for datasets with increasing number of genomes (Figure 4b) using a configuration mimicking a high-end desktop computer. Generating a database with all the optional analysis (except the RefSeq homologs search) took 1.9h, 3.9h, 8.6h, 21.0h and 55.6h for the datasets with 10, 20, 40, 90 and 179 genomes, respectively. The CPU time spent in the optional analyses increased linearly with the number of genomes This was however not the case for the core analysis. In particular, the cost of orthology prediction increased faster than the other analyses and will likely be the limiting factor for larger datasets. This is expected due to the *O*(*n*^2^) complexity of the all-against-all genomes comparison performed by Orthofinder (17).

**FIG 4.**
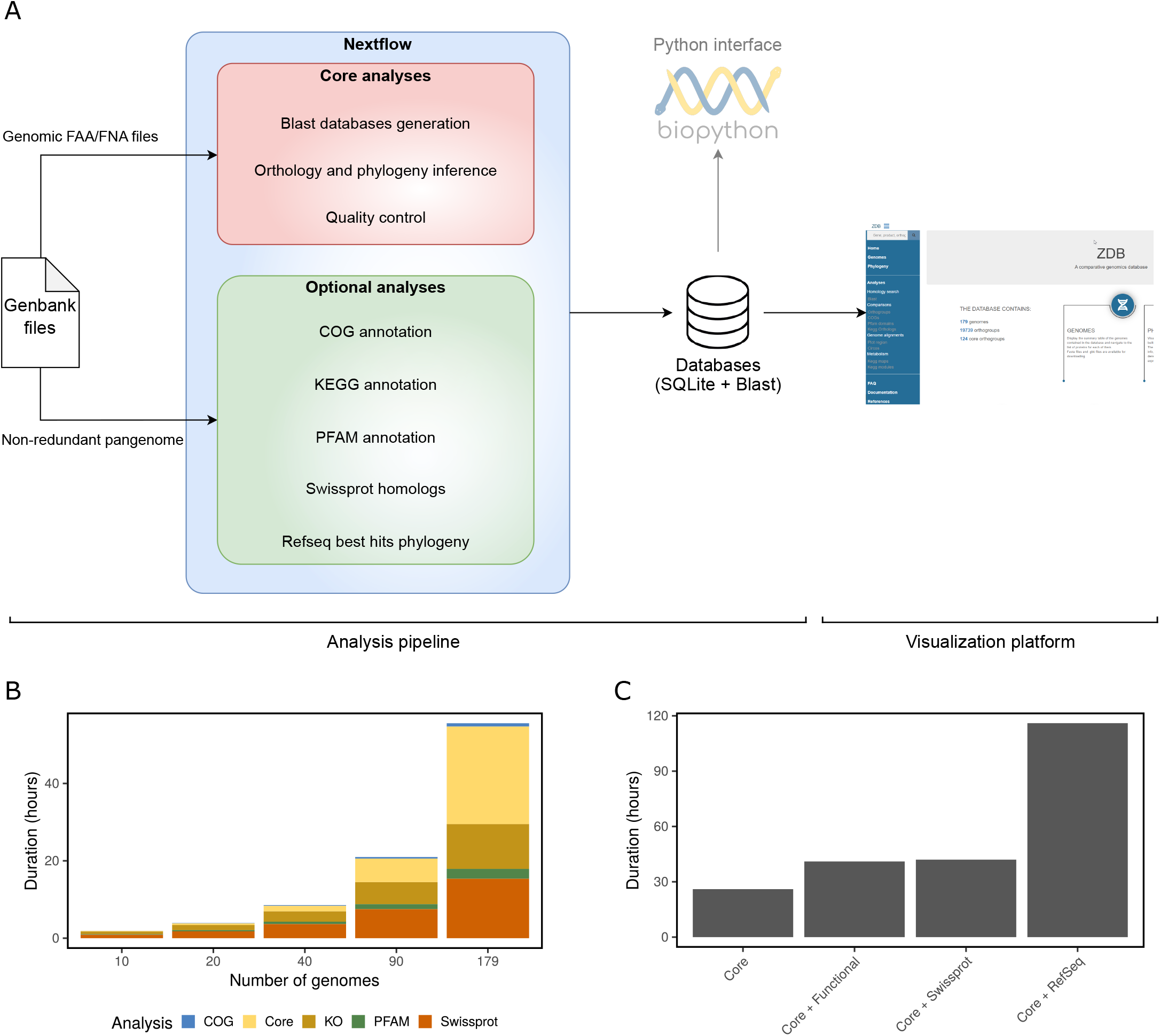
(A) zDB dataflow. The core analyses are performed for each dataset, while users can choose to perform additional optional analyses. (B) Duration of the analysis split by type (core and optional) in total CPU time according to the number of genomes in a benchmarking. (C) Benchmarking of the different analysis types with the 179 genome dataset. Functional analysis include COG and KEGG orthology annotations and Pfam domain prediction. Swissprot and RefSeq represent searching for homologs in Swissprot and Refseq, respectively.

The choice of optional analyses for the 179 genome dataset significantly impacts the computing time (Figure 4c). Despite the use of Diamond instead of blastp, searching for homologs in the RefSeq database took about 4 times longer than the other analyses. Similarly, searching for homologs in the Swissprot database took as long as performing the KEGG, COG and Pfam annotations together.

Finally, to confirm that zDB can process a hundred genomes on a mainstream computer, we ran the analysis pipeline on the 90 genomes dataset on a desktop machine, which took 71 hours to complete.

## DISCUSSION

zDB is a comparative genomics analysis tool entirely run on the user side that includes both an analysis pipeline and a web interface to visually explore the results. It was designed to require minimal typing on the command line; only three commands from installation to visualization of the results. As shown in the benchmarks, the analysis pipeline can process datasets of one hundred genomes in a matter of days on desktop computer, making dedicated computing infrastructures unnecessary for all but the largest datasets. The possibility to easily run Blast queries, to search for specific genes or to retrieve amino-acid and nucleotide sequences will make zDB useful for researchers more accustomed to lab work, while features such as core and pangenome analysis and genomic regions comparisons will be useful to seasoned bioinformaticians. The ability to share the database and launch the web interface from any computer facilitates data sharing and accessibility within and across groups and institutions with minimal infrastructure. Altogether, this makes of zDB a tool easy to use and install for a wide variety of applications such as genome browsing, characterization of newly sequenced genomes, or even setting up a public database for an organism of interest.

### Future directions

Large phylogenies are cumbersome to visualize as they may not entirely fit in a computer screen. To alleviate this, we plan to replace the ete3 drawing engine by a custom Javascript library to draw interactive phylogenetic trees allowing the user to collapse and expand branches. As of now, the addition or removal of genomes from an existing database is not possible and requires to repeat all the analysis on the modified dataset. We therefore plan to implement the possibility to add or remove genomes, which will allow users to incrementally improve a database without having to repeat the analyses. Finally, we plan to extend the set of optional analyses with additional annotations such as the prediction of antibiotic resistance genes, protein transmembrane domains and signal peptides. As the project is entirely open source and has been designed to be easily extended, the community is also welcome to implement any new features.

## MATERIALS AND METHODS

### Design and implementation

zDB is composed of two parts that can be run independently (Figure 4a): an analysis pipeline which performs all the computationally intensive steps and stores the results in a Sqlite3 database, and a visualization platform which renders the results stored in the database in a graphical interface.

The analyses are separated in a set of core analyses focused on orthology prediction and phylogeny inference and a set of independent optional analyses, with a focus on functional annotation. To simplify the installation and make the analyses reproducible and scalable, all steps are run either within docker (18) or singularity (19) containers or in conda environments, under the control of the Nextflow workflow manager (20). Nextflow allows the analyses to be easily scaled from high-performance clusters to desktop machines, while containers guarantee the reproducibility and ease of installation by packaging the tools in controlled environments. After the completion of the analysis pipeline, zDB can export the results as a compressed archive for subsequent use. The ability to export the results was developed to facilitate sharing and to accommodate the fact that the analysis may have to be run and exported from a high performance computing (HPC) cluster, where long term storage might not be possible due to disk space constraints.

The visualization platform was implemented as a Django website building upon the scaffold of ChlamDB (7). The Django server can either be instantiated on a desktop computer, for local access, or on an Internet-facing computer, if the website is to be made public (internally within a network or externally). The results are rendered as lists, annotated phylogenetic trees and interactive plots. The phylogenetic trees are drawn as static images with the ete3 toolkit (21), while the interactive plots are generated by a collection of home-made scripts based on the d3.js framework and several libraries such as jvenn.js (22), Circos.js (https://github.com/nicgirault/circosJS), BlasterJS (16) and plotly (https://plotly.com). All the plots drawn by the website can be downloaded as support vector graphics (.svg) images for subsequent use. Finally, users can also retrieve the results directly from the database via a Python (23) interface, if custom analyses are to be performed.

The code is available on GitHub (https://github.com/metagenlab/zDB) and zDB can be installed as a conda package.

### Minimal analyses: quality control, orthology inference and core genome phylogeny

The minimal set of analyses includes quality control with CheckM (24), the generation of Blast (25) databases, orthology prediction and phylogeny inference. This core analysis should be sufficient for most applications, as it allows core/pangenome analysis, gene annotation based on homology and generates species and gene phylogenies.

zDB takes GenBank files as input and has currently been tested with the output of Prokka (3), PGAP (26) and Bakta (27). As locally assembled genomes may have duplicated accessions or locus tags, zDB first checks their uniqueness and automatically generates new identifiers if necessary. Amino-acid and nucleotide sequences are then extracted from the GenBank files and used as input for subsequent analyses (4a). Annotations such as gene names and protein products are also extracted from the GenBank files. Those annotations are indeed particularly valuable when reference genomes are analyzed together with draft genomes, as the annotations of the genes from a reference genome may hint at the function of their homologs in draft genomes.

Orthology is inferred using Orthofinder (17). The sequences of orthologous proteins are aligned with MAFFT (28) and the alignments are used to infer phylogenetic trees for each orthogroup using FastTree (29). In addition to gene phylogenies, zDB also generates a species tree with FastTree using the concatenated alignments of the single-copy core orthologs. As some assemblies may be incomplete, the condition that core orthologs must be present in all genomes can be relaxed to allow missing genes. zDB generates Blast databases with both amino-acids and nucleotides sequences for each individual genomes and for the whole dataset. This allows users to search for sequences in a specific genome without the interference of better hits in other genomes, while still making it possible to perform global searches on the whole dataset.

### Optional analyses: homology search, COG, KEGG and Pfam annotations

To complement the core analysis, zDB can perform optional analyses focused on function prediction. Optional analyses all take the proteins of the non-redundant pan-genome as input and include the assignment to the Cluster of Orthologs Genes (COG), mapping to the Kyoto Encyclopedia of Genes and Genomes (KEGG), prediction of Pfam protein domains and search for homologs in the SwissProt database. COG annotations offer clues regarding protein functions and allow their classification in broad functional categories. The assignment to COG (37) clusters is performed by rps-blast (25) searches using the position-specific score matrices of the NCBI Conserved Domain Database (CDD) (35). KEGG annotations give insights into the metabolic capacities of the analyzed bacteria. The mapping to KEGG orthologs is performed by Kofamscan (34) using the prokaryotic profiles of the KEGG database (33). As they can offer functional insights into otherwise unannotated proteins and as domain architecture conservation may be a valuable addition to a gene phylogeny, Pfam protein annotations were also added in the optional analysis. The annotation is performed with the Pfam_scan (36) tool and the Pfam-A database. Finally, zDB can also perform homology search with blastp (25) against the manually curated entries of the SwissProt (30) database. The reference database used by zDB to perform those analysis are listed in Table 1. Of note, the core analyses can be performed without any reference database.

**TABLE 1.** Reference databases used by zDB

| Database | Release | Size <sup>a</sup> | Search tool |
| --- | --- | --- | --- |
| Swissprot (30) | Release 2021_04 | 86M | Blastp (25) |
| Refseq nr <sup>b</sup> (31) | Release 210 | 34.9G | Diamond (32) |
| KEGG hmm profiles (33) | Release 03/2022 | 1.2G | Kofamscan (34) |
| CDD (35) | Release 3.19 | 4.0G | Rpsblast (25) |
| Pfam-A hmm profiles (36) | Release 35.0 | 279M | Pfam_scan (36) |

<sup>a</sup>Refers to the volume of data to download
<sup>b</sup>As downloading the non-redundant RefSeq database is prone to failure, it is not automatically downloaded by zDB. A script to download and prepare the databases is installed with zDB, but has to be run manually.

To screen for lateral gene transfers using a well-validated method (38), zDB can search the RefSeq database for homologs of proteins from the non-redundant pangenome. The search is performed by Diamond (32) to reduce the duration of the analysis. The proteins of every orthogroup and their best hits (the best 4 hits of every protein, by default) are then aligned with MAFFT, and the alignment is used by FastTree to infer a phylogenetic tree. As reference genomes downloaded from RefSeq may have been included in zDB input dataset, the best hits from genomes already present in the database are filtered out. If the database was populated with genomes of related bacteria, observing that a protein from a distant taxa clusters more closely than the other proteins from the same orthogroup may indeed indicate a lateral gene transfer.

### Benchmarking

Although the analysis pipeline could process more genomes, the visualization platform is designed for datasets ranging from tens to hundreds of genomes. We therefore chose a representative dataset composed of the NCBI’s 179 reference genomes of the *Enterobacteriaceae* family to benchmark the analysis pipeline. The genomes were downloaded as Genbank files from the NCBI (the details are shown in supplementary Table 1). We ran a first benchmark to measure the running time of the different optional analyses on the full dataset. As the search for RefSeq homologs proved to be prohibitively long (Figure 4C), it was not included in the subsequent benchmark. The pipeline was then run on randomly generated subsets of the 179 genomes composed of 10, 20, 40, 90 or all genomes, all with a mean genome size of 3.8Mbp.

The performances of the pipeline were measured using Nextflow –with-report option. All analyses were run on an Ubuntu 18 server (112 Intel Xeon Platinum 8280 2.7GHz CPUs, equipped with 377GB of RAM memory), limiting parallelization to 20 simultaneous processes (with Nextflow cpus option) and total memory usage to 32GB (with Nextflow memory option), to mimic the computing power of a high-end desktop computer. We also tested the 90 genomes dataset on a desktop computer with 6 cores to have a better idea of the performances on a computer with more limited resources.

## SUPPLEMENTAL MATERIAL

**FIG S1**. The species tree generated by zDB, annotated with the main characteristics of the included genomes.

**TABLE S1**. Supplemental table 1 lists all the genomes used for the benchmarking of zDB.

## ACKNOWLEDGMENTS

The authors would like to thank the other members of the laboratory, Elena MontenegroBorbolla, Carmen Chen, Sedreh Nassirnia, Yangji Choi and Elindi De Coning, who tested the tool and provided feedback during the development. Bastian Marquis is funded by the Jürg-Tschopp MD-PhD scholarship. This work was supported as a part of NCCR Microbiomes, a National Centre of Competence in Research, funded by the Swiss National Science Foundation (grant number 180575).

